# Horizontal transfer and gene loss shaped the evolution of alpha-amylases in bilaterians

**DOI:** 10.1101/670273

**Authors:** Andrea Desiderato, Marcos Barbeitos, Clément Gilbert, Jean-Luc Da Lage

## Abstract

The subfamily GH13_1 of alpha-amylases is typical of Fungi, but it is also found in some unicellular eukaryotes (e.g. Amoebozoa, choanoflagellates) and non-bilaterian Metazoa. Since a previous study in 2007, GH13_1 amylases were considered ancestral to the Unikonts, including animals, except Bilateria, such that it was thought to have been lost in the ancestor of this clade. The only alpha-amylases known to be present in Bilateria so far belong to the GH13_15 and 24 subfamilies (commonly called bilaterian alpha-amylases) and were likely acquired by horizontal transfer from a proteobacterium. The taxonomic scope of Eukaryota genomes in databases has been greatly increased ever since 2007. We have surveyed GH13_1 sequences in recent data from ca. 1600 bilaterian species, 60 non-bilaterian animals and also in unicellular eukaryotes. As expected, we found a number of those sequences in non-bilaterians: Anthozoa (Cnidaria) and in sponges, confirming the previous observations, but none in jellyfishes and in Ctenophora. Our main and unexpected finding is that such fungal (also called Dictyo-type) amylases were also consistently retrieved in several bilaterian phyla: hemichordates (deuterostomes), brachiopods and related phyla, some molluscs and some annelids (protostomes). We discuss evolutionary hypotheses possibly explaining the scattered distribution of GH13_1 across bilaterians, namely, the retention of the ancestral gene in those phyla only and/or horizontal transfers from non-bilaterian donors.

## Introduction

Alpha-amylases are enzymes that are almost ubiquitous in the living world, where they perform the hydrolysis of starch and related polysaccharides into smaller molecules, to supply energy to the organism through digestion. They belong to glycosyl hydrolases, a very large group of enzymes which have been classified in a number of families according to their structures, sequences, catalytic activities and catalytic mechanisms (Henrissat and Davies 1997). Most alpha-amylases are members of the glycoside hydrolase family 13 (GH13), which includes enzymes that can either break down or synthetize α −1,4-, α −1,6- and, less commonly, α −1,2- and α −1,3-glycosidic linkages. Sucrose and trehalose are also substrates for enzymes of this family (MacGregor *et al.* 2001). The numerous family GH13 is divided into 42 subfamilies, of which only three occur in Metazoans: GH13_1, GH13_15 and GH13_24 (Stam *et al.* 2006; Da Lage *et al.* 2007; Lombard *et al.* 2014). The latter two include the common animal alpha-amylases, while the former was first described in Fungi for which it represents the canonical alpha-amylase (Stam *et al.* 2006). Da Lage *et al.* (2007) described the subfamilies GH13_15/24 as private to Bilateria among metazoans. In the same article, they retrieved sequences belonging to the subfamily GH13_1 from the sponge *Amphimedon queenslandica* (named *Reniera sp.* in their paper) and the sea anemone *Nematostella vectensis,* besides the unikont choanoflagellates and amoebozoans, and also excavates and ciliates. They dubbed “Dictyo-type” this alpha-amylase, referring to the slime mold *Dictyostelium discoideum* (Amoebozoa Mycetozoa). The authors proposed that this amylase, ancestral to the Unikont clade, is shared among non-bilaterian metazoans (e.g. sponges, sea anemones and corals, and Placozoa), but was replaced in Bilateria by an alpha-amylase of bacterial origin, whose sequence is close to the typical animal amylases.

Given that a wealth of new genomes have been sequenced in the twelve years after that publication, we decided to explore again the diversification of this enzyme subfamily among the Eukaryota. We will focus mainly on Metazoa, in which we show unexpected situations of cooccurrence of both subfamilies GH13_1 and GH13_15/24 in the same genomes. We will discuss two mutually exclusive explanations that may be proposed: either the retention of the ancestral GH13_1 gene along with the typical bilaterian GH13_15/24 in some phyla, or horizontal transfer(s) from non-bilaterian animal donor(s) which would have to be identified.

## Materials and methods

In order to further characterize the distribution of GH13_1 genes in Metazoa, we used the sequence of the sponge *Amphimedon queenslandica* GH13_1 (GenBank XP_019851448) as a query to perform BLASTP and TBLASTN searches on various online databases available in Genbank (nr, proteins, genomes, assembly, SRA, TSA, WGS), and also in the more specialized databases compagen.org, marinegenomics.oist.jp, reefgenomics.org, marimba.obs-vlfr.fr, vectorbase.org, PdumBase (pdumbase.gdcb.iastate.edu), AmpuBase (https://www.comp.hkbu.edu.hk/~db/AmpuBase/index.php) (Ip *et al.* 2018), between October 2018 and August 2019. Fungi were not searched further in this study because they are known to have a GH13_1 member as the usual alpha-amylase. To increase the chances to retrieve potential cnidarian or ctenophoran sequences, the starlet sea anemone *Nematostella vectensis* amylase (XP_001629956) was also used to query those databases. After the discovery of GH13_1-like sequences in Bilateria, the sequence XP_013396432 of the brachiopod *Lingula anatina* was also used for specific search in Bilateria. Non-animal eukaryote species were investigated using the *Dictyostelium discoideum* sequence XP_640516 as query. We chose a stringent BLAST threshold because glycosyl hydrolases from other GH13 subfamilies might be retrieved otherwise, owing to the presence of stretches of amino acids that are conserved across the subfamilies despite other enzymatic specificities (Janeček 1994; Janeček *et al.* 2014; MacGregor *et al.* 2001; van der Kaaij *et al.* 2007). Therefore, the BLAST hits (or High-scoring segment pairs HSPs) were considered further when expectation values (e-values) were better (lower) than 1e-100 for BLASTP or 1e-75 for TBLASTN in annotated or assembled genomes or transcriptomes which returned full-size or near full-size GH13_1 sequences. When only partial sequences could be retrieved using BLAST we collected a large genome region encompassing and flanking the BLAST hit and tried to reconstitute the full-size sequence. The stringent threshold was obviously not applied to constitutively small HSPs such as SRA (sequence read archives). These were only considered when several highly significant hits (typically 1e-10) covered a large part of the query sequence. Since SRA HSPs generally did not overlap, we could not assemble longer sequences and thus we did not use such sequences in alignments or phylogenies. SRA and transcriptome databases are prone to contamination, thus we checked by reciprocal BLAST that the retrieved sequences were not contaminations. SRA databases were used firstly when no or few assembled genomes or transcriptomes were available (e.g. Nemertea, Bryozoa). If a GH13_1 sequence was found in an annotated or assembled genome, SRA HSPs were also used in order to increase the sampling of related taxa and then added some support to the presence of GH13_1 in the lineage considered. On the other hand, we considered that within a given lineage, the absence of GH13_1 sequence in a reliable annotated or assembled genome, combined to the detection of GH13_1 in SRA databases from related species would suggest that the GH13_1 gene was lost within this lineage in some taxa but not all. Conversely, if no GH13_1 sequence at all was found in any annotated genome and in any other database, we considered that the gene was lost in an ancestor of this lineage. When working with “assembled genomes” (non-annotated), we reconstituted exon-intron gene structure as well as the the protein sequence from the TBLASTN results. Finally, for phylogenetic analyses we kept only sequences which lay inside long contigs, or full-size or near full-size transcripts. We also checked once again the absence of animal-type alpha-amylase (GH13_15 or 24) outside the Bilateria using the sequence of the bivalve *Corbicula fluminea* (AAO17927) as a BLASTP query. The CAZy database (cazy.org (Lombard *et al.* 2014)), which is devoted to glycosyl hydrolases and related enzymes was used to check assignment of some of the sequences we found to the GH13_1 subfamily.

Intron-exon gene structures were recovered either from alignments between genomic sequences and their mRNA counterparts, or using annotated graphic views when available in the databases. In cases of likely erroneous annotations we reanalyzed the gene region by eye, correcting dubious frameshifts if necessary (see. Fig. S1 as an example). In some cases, for unannotated genes, the N-terminal and/or the C-terminal parts of the retrieved genomic sequences were uncertain, and were not retained in the analyses.

Alignments were performed using MUSCLE (Edgar 2004), as implemented in Geneious (Biomatters Ltd.). A maximum likelihood (ML) tree was built using PhyML’s (Guindon and Gascuel 2003) current implementation at the phylogeny.fr portal (Dereeper *et al.* 2008). To this end, we first trimmed the N-terminal protein sequences up to the first well conserved motif LLTDR. C-terminal parts were also truncated at the last well aligned stretch. Gaps were removed from the alignments and data were analyzed under WAG (Whelan and Goldman 2001) with among-site rate variation modeled by four discretized rate categories sampled from a gamma distribution. Both the alpha parameter and the proportion of invariable sites were estimated from the data. The robustness of the nodes was estimated using an approximate likelihood ratio test (aLRT) (Anisimova and Gascuel 2006). The tree was drawn at the iTOL website (Letunic and Bork 2016). Metazoans and choanoflagellates were clustered as the ingroup.

The protein sequences, Fasta alignment and Newick-formatted tree are available at FigShare: https://figshare.com/articles/GH13_1_metazoa/9959369. Supplementary Figures and Tables are available at FigShare: https://figshare.com/articles/Suppl_data_GH13_1/9975956

## Results

The sequences retrieved from the databases are listed in Table 1. The metazoans investigated are listed in Tables S1 (non-bilaterians) and S2 (bilaterians) with indication of the current state of genome/transcriptome sequencing, the database, the presence or absence of GH13_1 sequences, and the number of gene copies, where possible. A general protein alignment of the sequences found in this study along with already known GH13_1 sequences is shown in Fig. S2.

**Table 1:**
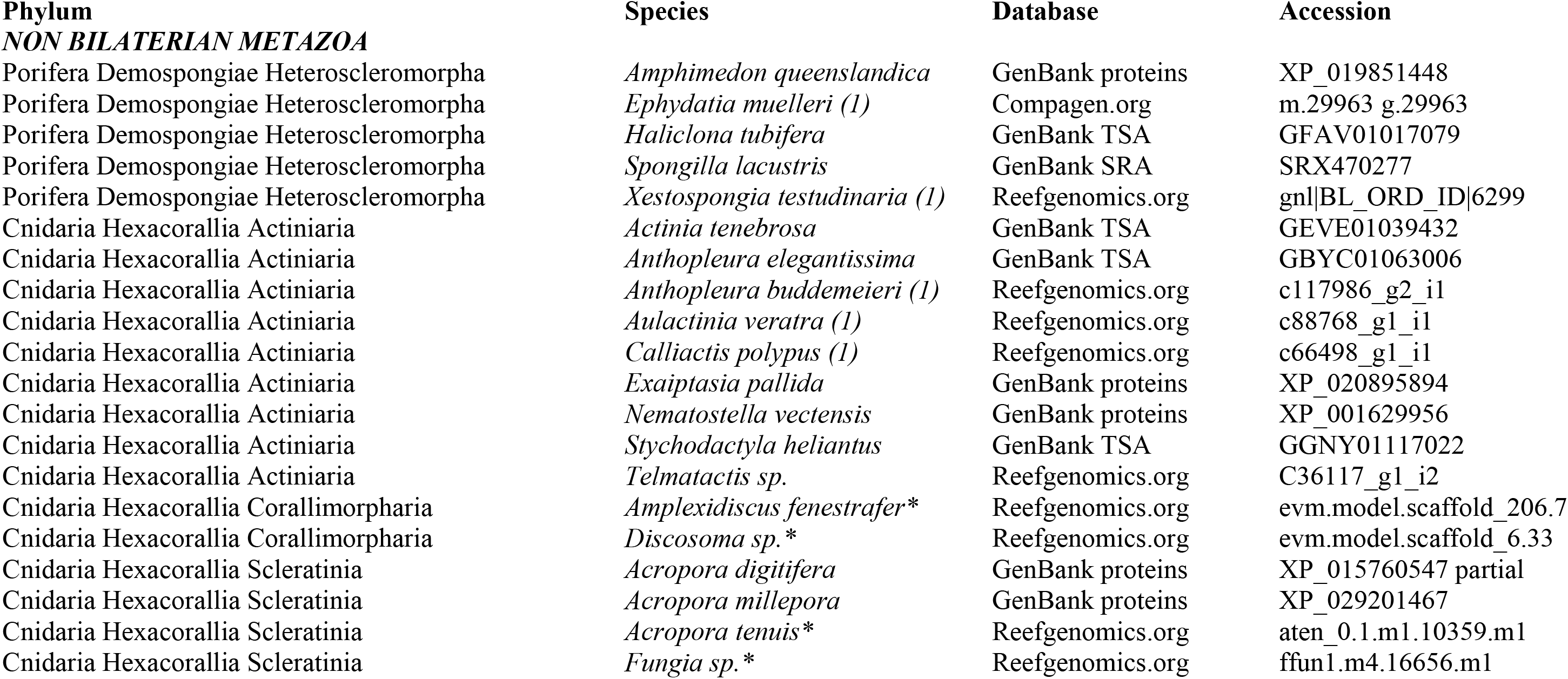
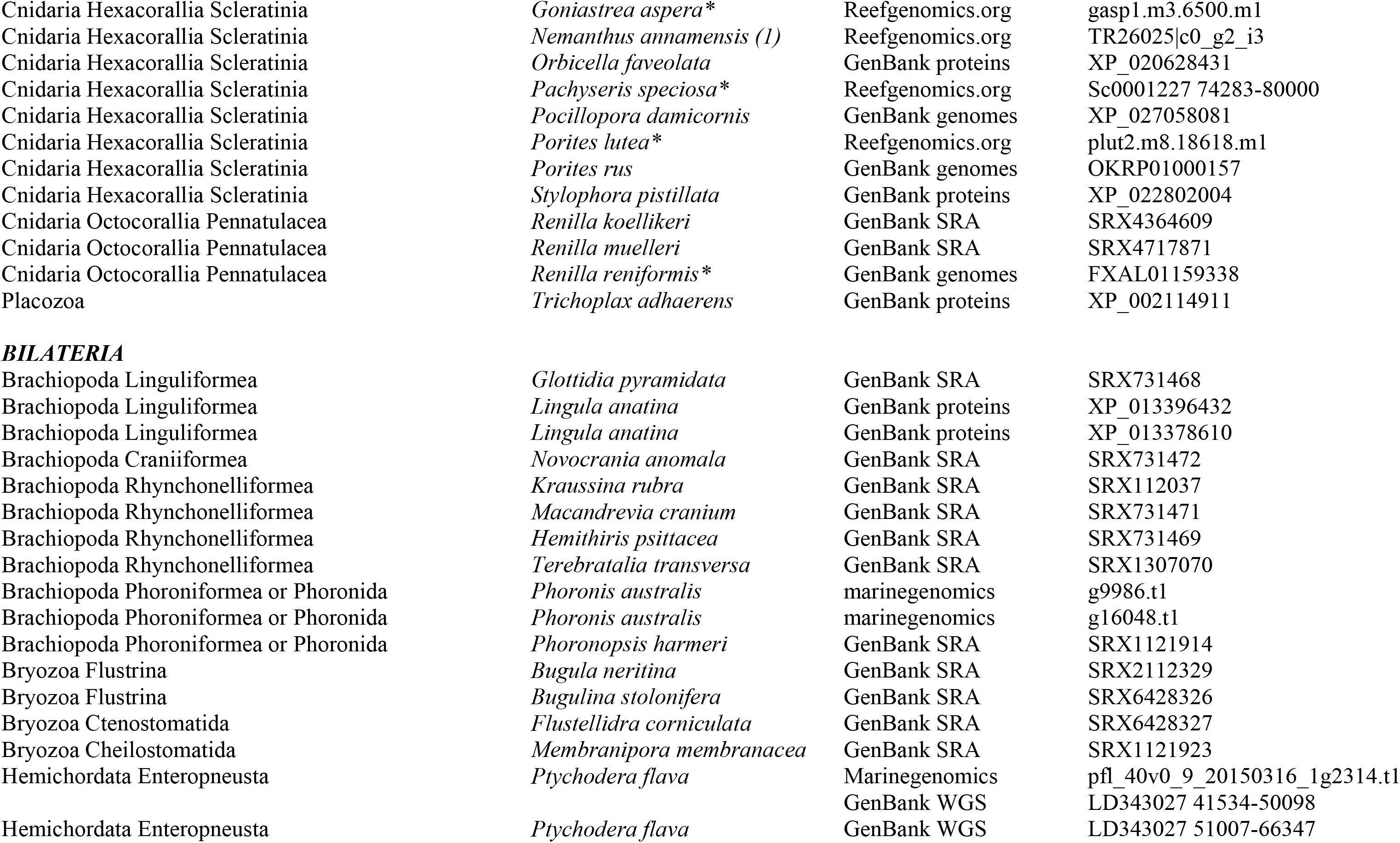
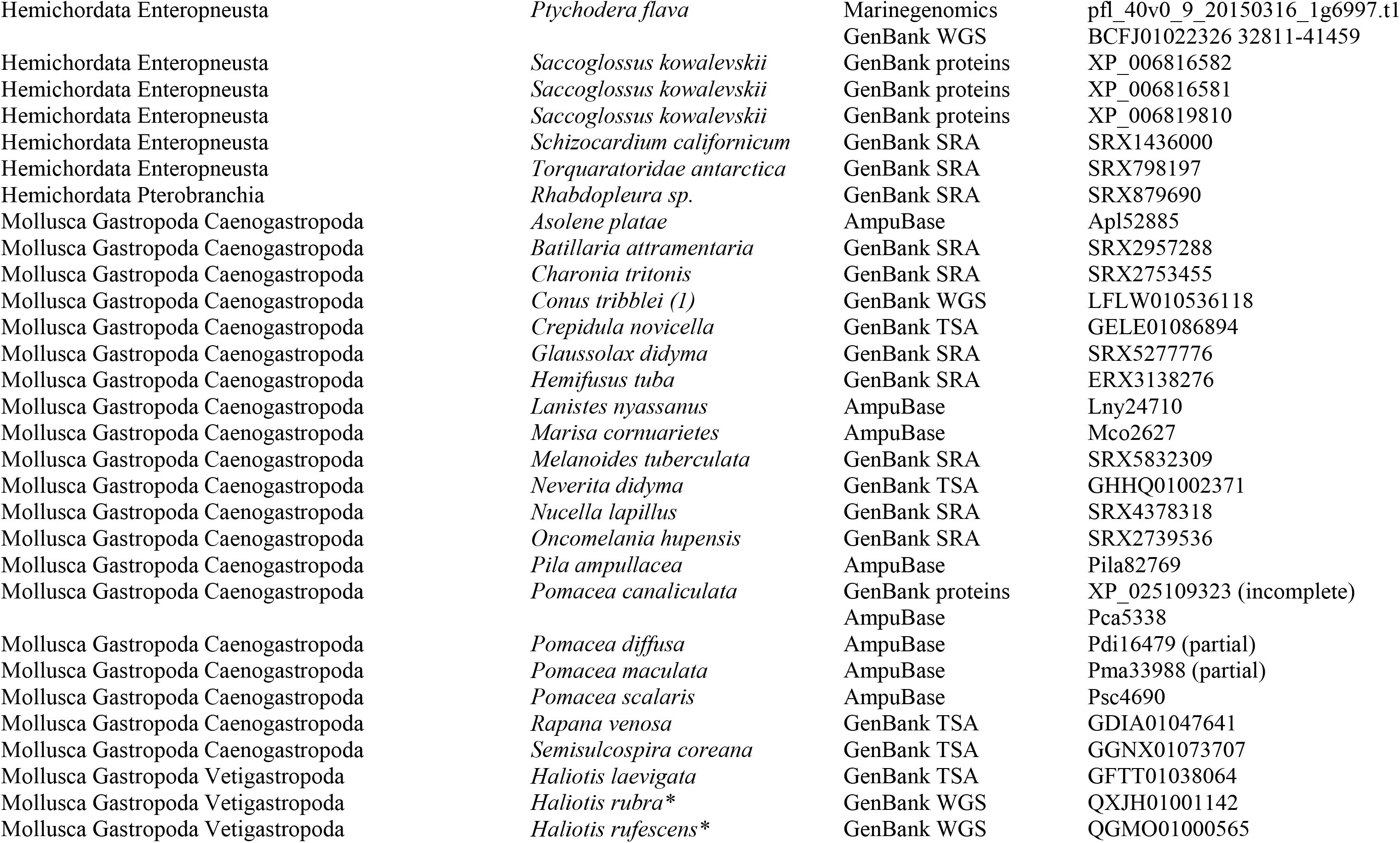
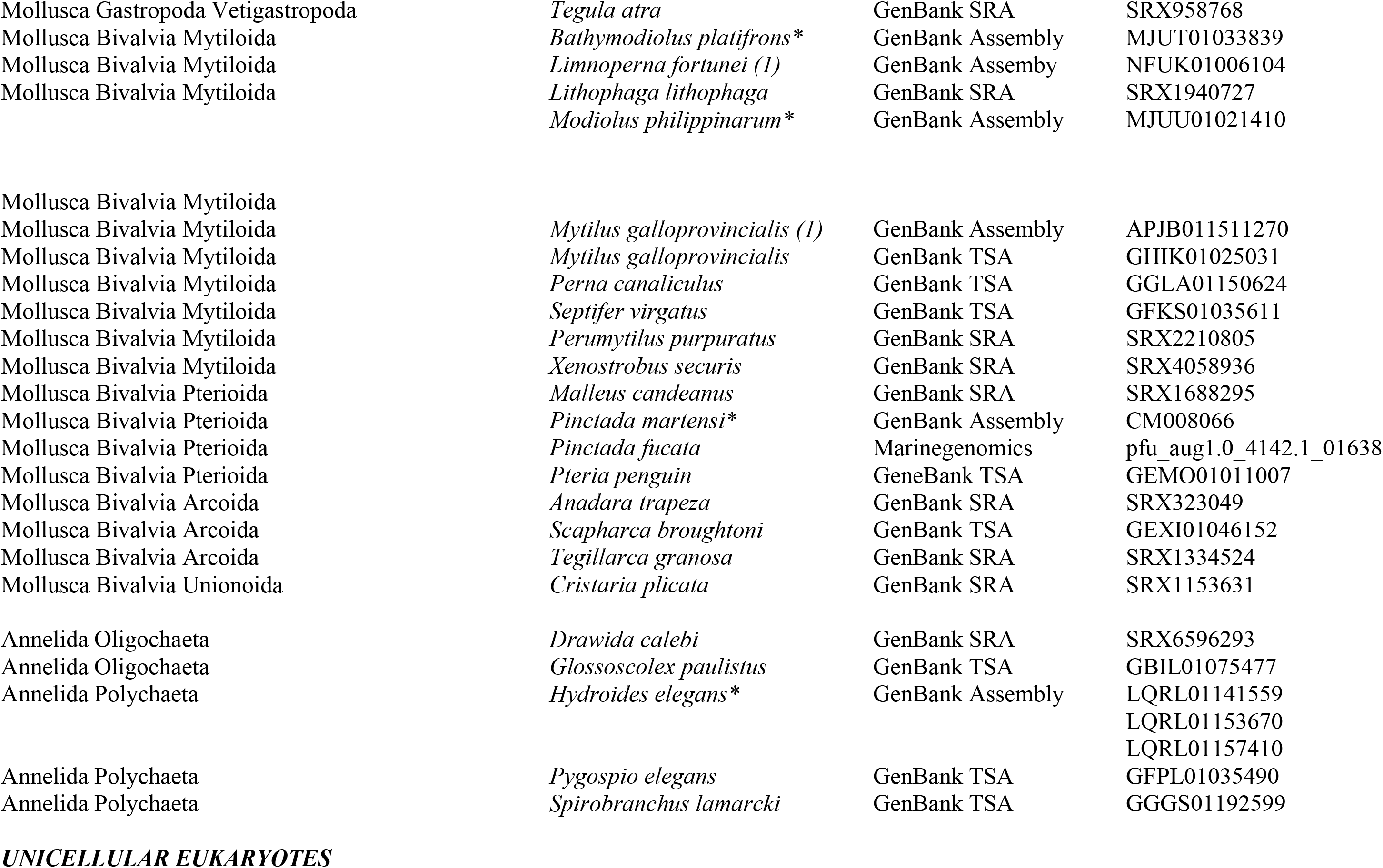
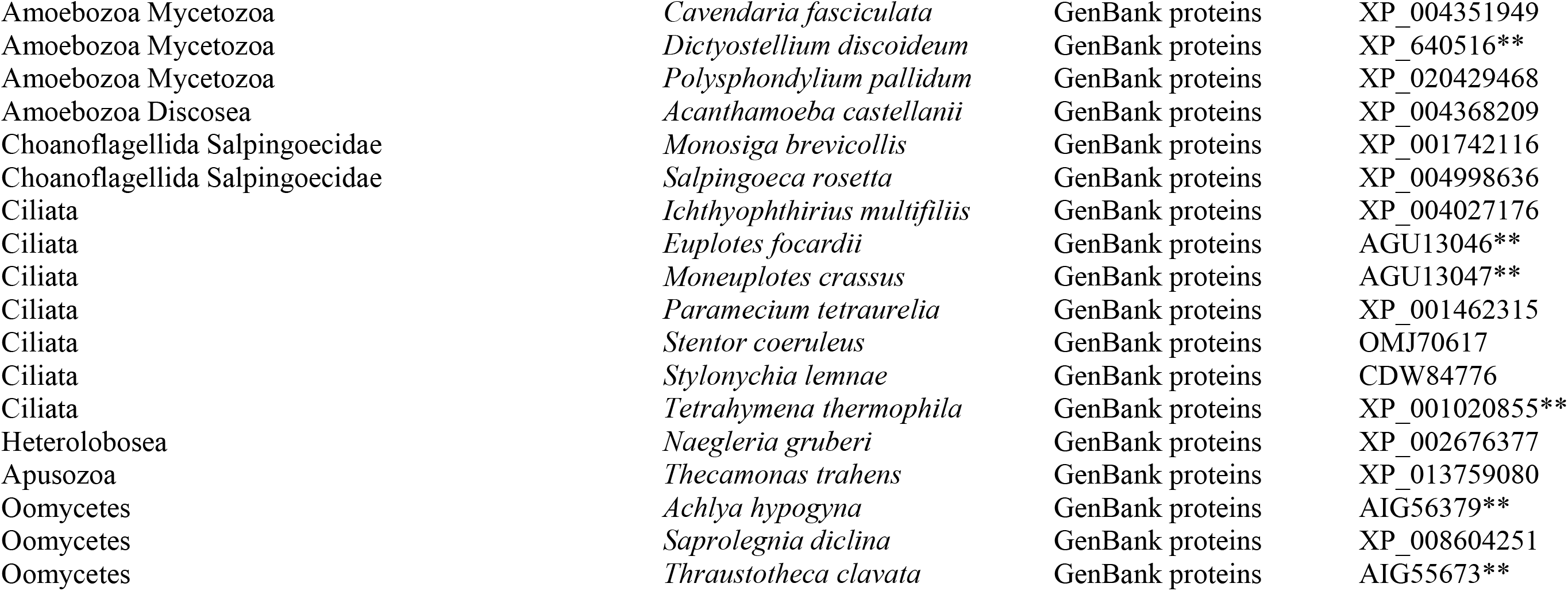
GH13_1-like sequences found after BLAST searches in online databases (not comprehensive for unicellars, without the Fungi). *: sequences which have not been characterized as protein-coding, in sequenced genomes with long contigs; (1): from short DNA sequences (except Sequence reads archive); **: reported as GH13_1 in CAZy. Most of the SRA data are from transcriptome studies; see Tables S1 and S2.

### GH13_1 sequences retrieved from unicellular taxa

We confirmed the presence of GH13_1 in dictyostelids, in ciliates and also in oomycetes, some representatives of which (but not all) are indicated in Table 1. In two oomycetes, *Saprolegnia diclina* and *Achlya hypogyna,* the GH13_1-like sequences were the C-terminal half of longer sequences, the N-terminal half of which was similar to unclassified GH13 sequences found in e.g. *Acanthamoeba histolytica* (GenBank accession BAN39582), according to the CAZy database. In our general phylogenetic tree (Fig. 1), these sequences were used as outgroups. In choanoflagellates, where *Monosiga brevicollis* was already known to harbor a GH13_1 sequence (Da Lage *et al.* 2007), we found a GH13_1 sequence in the genome of *Salpingoeca rosetta.* A partial sequence was also returned from incomplete genome data from *Monosiga ovata* (at Compagen, not shown).

**Figure 1:**
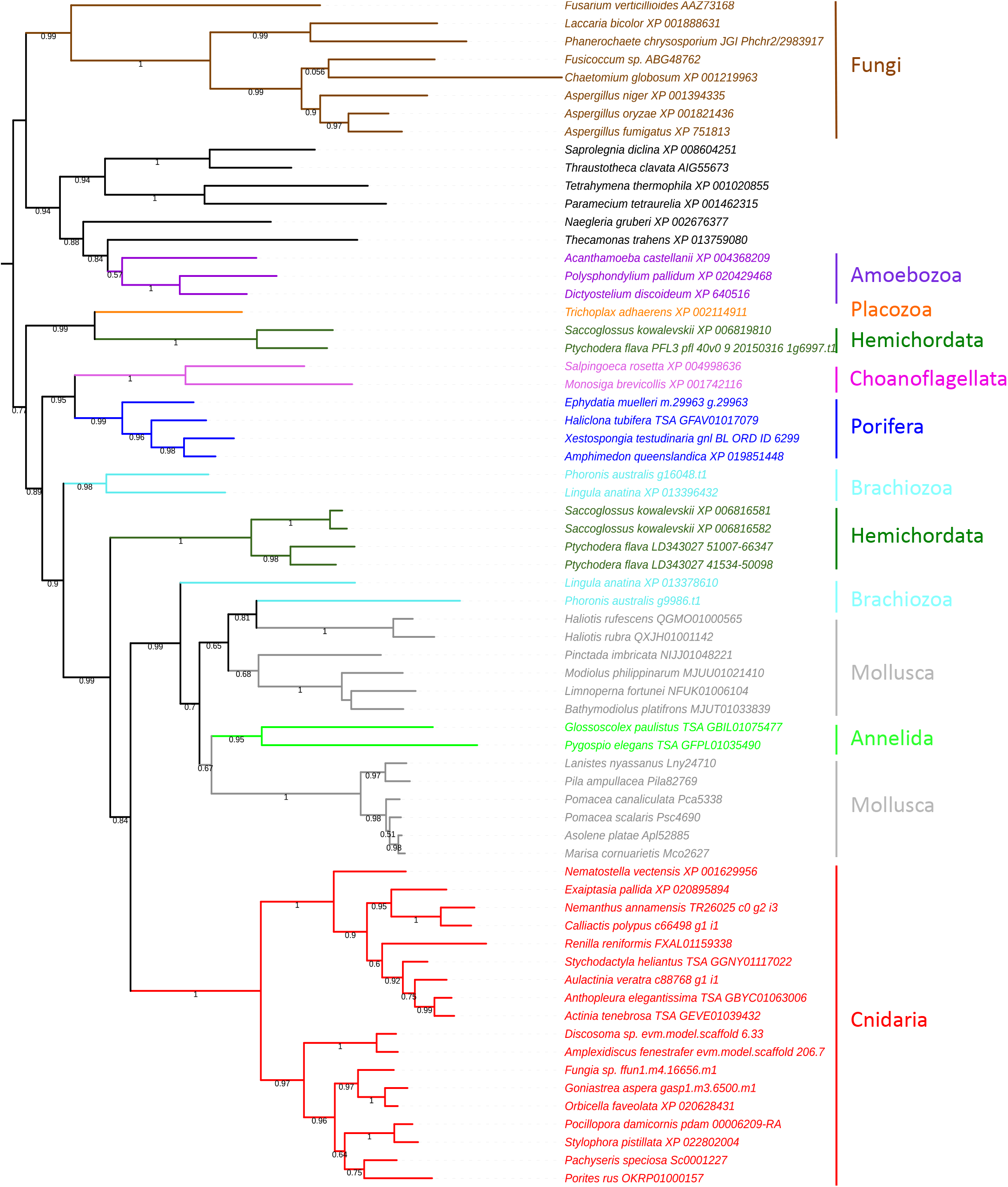
ML tree of GH13_1 protein sequences of metazoan and non-metazoan species. The tree was rooted by placing fungi and unicellular organisms, except choanoflagellates, as outgroups. The numbers at the nodes are the aLRT supports. Dark green: hemichordates; light blue: brachiozoans; red: cnidarians, dark blue: sponges; orange: placozoans; pink: choanoflagellates; purple: amoebozoans; brown: fungi; grey, molluscs; bright green: annelids; black: other protists.

### GH13_1 sequences retrieved from non-bilaterian animals

In Cnidaria, a number of GH13_1 sequences were recovered from many Anthozoa species (sea anemones, corals and allies), from genome as well as transcriptome data, at the Reefgenomics database (Table S1). Interestingly, we found no alpha-amylase sequences at all in Medusozoa (jellyfishes, hydras) nor in Endocnidozoa (parasitic cnidarians). In the general tree (Fig. 1), cnidarian sequences form a clear cluster with two main branches, grouping Actiniaria (sea anemones) and Pennatulacea (soft corals) on one branch, and Scleractinia (hard corals) and Corallimorpharia (mushroom anemones) on the other branch.

In sponges (Porifera), data were less abundant. No alpha-amylase sequence was found in *Sycon ciliatum* (Calcarea) and *Oscarella carmela* (Homoscleromorpha). All the sequences we retrieved belonged to Demospongiae. Similarly, we found no amylase sequence at all in the phylum Ctenophora (*Mnemiopsis leidyi, Pleurobrachia bachei*), the phylogenetic position of which is controversial: it has been recovered as the most basal metazoan (Whelan *et al.* 2017), as Cnidaria’s sister group e.g. Simion *et al.* 2017, Philippe *et al.* 2009), re-establishing Coelenterata, and also as the earliest branch in the Eumetazoa (animals with a digestive cavity and/or extra cellular digestion) e.g. Pisani *et al.* 2015.

### GH131 sequences retrieved from bilaterian animals

The surprising finding of this study, on which we will focus our attention, is the consistent, albeit sparse, occurrence of GH13_1 alpha-amylase sequences in several bilaterian phyla: hemichordates, which are deuterostomes, brachiopods, phoronids (Brachiozoa) and Bryozoa, and in some molluscs and annelids (Eutrochozoa), which are all protostomes. In the well annotated genomes of the brachiopod *Lingula anatina* and the phoronid *Phoronis australis*, two paralogs were found (Table 1). In both species, the two copies are located on different contigs. The paralog sequences are rather divergent, given their positions in the tree (Fig 1) and each paralog groups the two species together. This indicates that not only duplication, but also the divergence between paralogs is ancestral to these species, dating back at least to basal Cambrian, according to the TimeTree database (Kumar *et al.* 2017). GH13_1 sequences were found in other brachiopods and phoronids as sequence reads (SRA) from transcriptome data only, with no available genomic support (listed in Table 1 and S2). We must be cautious when only transcriptome data are available, as transcripts from contaminating symbionts or parasites may generate false positives (Borner and Burmester 2017) and/or the lack of expression of the targeted sequence in the investigated tissues may lead to false negatives. However, seven different brachiopod species returned positive hits, giving some robustness to our finding. Two phyla are related to Brachiozoa: Bryozoa and Nemertea (Kocot 2015; Luo *et al.* 2018, but see Marlétaz *et al.* 2019). We found clues for the presence of GH13_1 in four Bryozoa species, but only transcriptome reads were available. In contrast, in Nemertea, none of the 14 species investigated returned any GH13_1 sequence, including the annotated genome of *Notosperma geniculatus*.

Similarly, we found three gene copies in the genomes of the hemichordates *Saccoglossus kowalevskii* and *Ptychodera flava.* In both species, two copies are close to each other (XP_006816581 and XP_006816582 in *S. kowalevskii,* and their counterparts in *P. flava)* as shown by the topology of the gene tree (Fig. 1). This could suggest independent gene duplication in each species. However, we observed that the two duplicates were arranged in tandem in both species, which would rather suggest concerted evolution of two shared copies. In *P. flava,* this genome region is erroneously annotated as a single gene at the OIST Marine Genomics database. The third paralog is very divergent from the two other copies, so its divergence from the ancestral copy probably occurred before the species split, as well. The three copies were therefore probably already present before the split of the two lineages, some 435 mya (Kumar *et al.* 2017). Three other hemichordate species, *Schizocardium californicum, Torquaratoridae antarctica* and *Rhabdopleura sp.* harbor a GH13_1 gene, as shown by SRA search in GenBank (Table 1). A positive result was also retrieved from the genome of *Glandiceps talaboti* (Héctor Escrivà, Oceanology Observatory at Banyuls-sur-mer, personal communication).

In molluscs, we found BLAST hits with significant e-values in gastropod species from two clades only, the Vetigastropoda (e.g. the abalone *Haliotis* sp.) and the Caenogastropoda (e.g. Ampullariidae such as *Pomacea canaliculata).* We consistently found one copy in eight species belonging to the family Ampullariidae. In *P. canaliculata,* the genome of which has been well annotated, the GH13_1 sequence (XP_025109323) lies well inside a 26 Mbp long scaffold (linkage group 10, NC_037599) and is surrounded by *bona fide* molluscan genes (Table S3). GH13_1 sequences were found in other Caenogastropoda from SRA or transcriptome databases (Table 1 and S2). We also found GH13_1 sequences in several bivalve clades: Mytiloida (e.g. the mussel *Mytilus galloprovincialis),* Pterioida (e.g. the pearl oyster *Pinctada imbricata),* Arcoida (e.g. *Scapharca broughtoni)* and in the Unionoida *Cristaria plicata.* For sequences retrieved from the TSA or SRA databases (see Table 1), whose issues were mentioned above, we performed reciprocal BLAST in GenBank nr. Almost always *Lingula anatina* was recovered as the best hit. However, as an example of the necessary careful examination of results, we found a significant HSP in a transcriptome database of the sea hare *Aplysia californica* (TSA GBDA01069500). This sequence was not found in the well annotated *A. californica* genome, and turned out to be related to ciliates. We found no occurrence of GH13_1 in Veneroida, Pectinoida and Ostreoida, for which annotated and/or assembled genomes exist, nor in cephalopods.

In annelids, we found occurrences of GH13_1 genes in a few species, the genomes of which are still not fully assembled, namely the “polychaetes” *Hydroides elegans, Pygospio elegans* and *Spirobranchus lamarcki* but not in the well-annotated genome of *Capitella teleta.* We also recovered HSPs from the clitellate *Glossoscolex paulistus* but not from *Amynthas corticis* or *Eisenia fetida.* We found no GH13_1 sequences in Hirudinea (leeches). To summarize, in molluscs as well as in annelids, the presence of GH13_1 genes is scattered and patchy across and within lineages. Interestingly, we found that some of the mollusc GH13_1-like sequences, especially in bivalves, were much shorter, either truncated at the C-terminal, or this region was so divergent from the query sequence *(L. anatina)* that it was impossible to identify, assemble and align it with our data set (Fig. S2). In addition, we found that the annelid *Hydroides elegans* had an internal deletion, which precluded its inclusion in the phylogenetic analysis. This suggests that those sequences may not have alpha-amylase activity.

### Gene tree analysis: position of bilaterian sequences

The goal of the gene tree analysis is to examine whether the occurrence of GH13_1 genes in bilaterian animals may be due to independent horizontal gene transfers (HGT) or if they descend from a GH13_1 alpha-amylase copy ancestral to Unikonts. In the first case, the bilaterians GH13_1 sequences are unlikely to cluster together and the gene tree topology will likely display one or more nodes that are inconsistent with the bilaterian phylogeny. In the second case, the bilaterian sequences are expected to recover a bilaterian clade and to have a cnidarian clade as its sister group (Laumer *et al.* 2018). The actual tree topology (Fig. 1) is not that straightforward when it comes to the bilaterian relationships, although we may rule out any proximity of bilaterians GH13_1 sequences with unicellular or fungal sequences, regardless of tree rooting.

All Cnidarian orthologs form a well-supported cluster. The sister relationship between Corallimorpharia and Scleractinia reflects what was recovered in species trees using different markers (e.g. Rodríguez *et al.* 2014), although the Scleractinia topology disagrees with previous phylogenetic analyses of the order (e.g. Barbeitos *et al.* 2010). The other cluster within Cnidaria is mainly composed of actiniarian (sea anemones) sequences, but it also includes, with strong support, the sequence queried from the sea pen *Renilla reniformis* (order Pennatulacea). This order belongs to the sub-class Octocorallia and not to Hexacorallia, the monophyletic sub-class in which scleractinians, corallimorpharians and sea anemones are found (e.g. Rodríguez *et al.* 2014). We used RAXML-NG v0.80 (Kozlov *et al.* 2019) to conduct a constrained search under WAG for a ML tree in which Hexacorallia was monophyletic and *R. reniformis* was placed as its sister group (e.g. Chang *et al.* 2015; Zapata *et al.* 2015)) and employed a simple LR test to statistically evaluate the difference betweeen the observed and expected (phylogenetic) placement of the *R. reniformis* sequence (Kozlov *et al.* 2019). The log-likelihood difference between the unconstrained (lnLh = −29,155.37) and constrained (lnLh = −29,208.38) ML tree scores was 53.01. According to Kass and Raftery (1995), there is very strong support for the highest likelihood hypothesis (in our case, the ML tree in Fig. 1) when the double of this difference (i.e. 2 x 53.01 = 106.02) exceeds 10 log-likelihood units. Thus, there is significant inconsistency between the position of *R. reniformis*’ GH13_1 copy and the phylogenetic placement of this species. This may be due to a horizontal transfer event that would have occurred within Cnidaria, but additional data from well-sequenced Pennatulacea would be welcome to check this possibility. Nevertheless, it is noteworthy that the genome of *Dendronephthya gigantea* (Octocorallia, order Alcyonacea) returned no result. Most bilaterian sequences are clustered with Cnidaria, as phylogenetically expected in the case of a shared ancestral gene, as a robust cluster grouping one Brachiozoa (brachiopod/phoronid) copy, the molluscs and the annelids, which is consistent with the phylogeny. However, the tandem hemichordate duplicates and the other Brachiozoa genes are not included in the bilaterian clade, but remain ingroup relative to the sponge sequences.

Interestingly, the two remaining hemichordate sequences are the earliest diverging lineage of the Metazoa + Choanoflagellata cluster, since they are branched with the placozoan *Trichoplax adhaerens* sequence, this relationship being strongly supported whatever the tree reconstruction method employed (Fig. 1, and data not shown). In order to check for the possibility of a long branch attraction (LBA), which would artificially cluster hemichordate and placozoan sequences, we performed Tajima’s relative rate tests (Tajima 1993) using MEGA7 (Kumar *et al.* 2016). The sequence of *S. kowalevskii* XP_006819810, suspected to evolve fast, was compared with its paralog XP_006816581, using five different outgroups, i.e. the three sponges and the two choanoflagellates. Unexpectedly, the χ^2^ tests returned non-significant values in two tests and significant values in three tests (Table S4). Therefore, with our data, LBA cannot be entirely ruled out in this particular case.

### Analysis of intron positions

Intron positions may be valuable markers when reconstituting gene histories. We identified 56 intron positions from the subset of species of the general tree for which we could find data (Fig. 2). Only one intron position is widely shared among these GH13_1 gene sequences. It is the first position reported in the alignment, and it lies just upstream to the first conserved part of the alignment. The main observation is the numerous conserved positions across bilaterian sequences (10 positions), and between bilaterian sequences and the sponge and the Placozoa (7 positions). In addition, three positions are common to bilaterians and the choanoflagellate *Monosiga brevicollis.* In contrast, the Cnidaria have few introns, with positions different from the sponge and the bilaterians, except for position 1. The other species under examination, i.e. protists and fungi, have essentially specific intron positions. Therefore, the overall conservation of intron positions across bilaterians + sponges is a further argument to state that an explanation of the occurrence of GH13_1 alpha-amylases in some bilaterians does not involve non-animal species.

**Figure 2:**
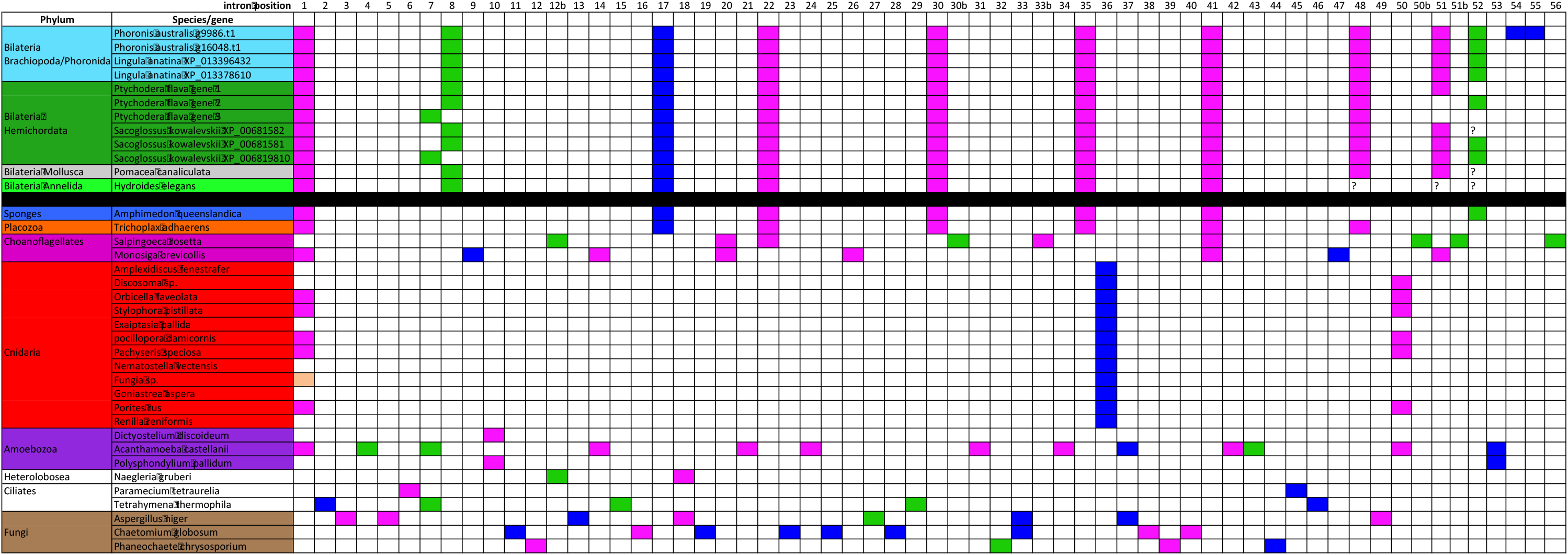
Intron positions compared across the sampled GH13_1 genes. The intron positions found in the studied parts of the sequences were numbered from 1 to 56. Pink: phase zero introns; green: phase 1 introns; blue: phase 2 introns. The black horizontal bar separates bilaterians from species where GH13_1 alpha-amylases are considered native. The color code for species is the same as in Figure 1.

## Discussion

The evolutionary scenario proposed by Da Lage *et al.* 2007, suggested that the GH13_1 alpha-amylase gene ancestral to Unikonts (Amoebozoa and Opisthokonts, i.e. Fungi and Metazoa/Choanoflagellata) was totally absent from Bilateria, due to its complete replacement by a new alpha-amylase, originating from a bacterium through HGT. Here, we have shown that a limited number of bilaterian lineages, all aquatic species, namely hemichordates, brachiozoans, bryozoans, and some sparse molluscs and annelids, actually do harbor GH13_1 alpha-amylase genes. Note that all those species also have at least one classical animal alpha-amylase of the GH13_15/24 subfamilies. Several species with whole genome well sequenced and annotated were found to harbor such genes in each phylum Hemichordata, Brachiozoa and molluscs. They were investigated in more details, especially regarding the genomic environment of their GH13_1 genes. We are quite confident that the GH13_1 sequences we found are not due to contaminating DNA. First, the bilaterian sequences retrieved from annotated genomes were inside long contigs, and mostly surrounded by genes showing bilaterian best BLAST hits (Table S3). However, the *S. kowalevskii* XP_006819810 gene could appear somewhat dubious, since it is placed at the distal end of a contig, with only two other genes on the contig (Table S3), one of which has a placozoan best hit. But its *P. flava* counterpart is well inside a gene-rich contig. Therefore, these seemingly non-bilaterian genes are well in bilaterian genomic contexts. Second, a lot of additional sequences from other species belonging to these phyla were gathered from more sketchy data, i.e. lower-quality assembled genomes, transcriptomes or sequence read archive databases, which added some support to the presence of these amylase genes. Although transcriptome and rough genomic data should be handled with care, this lends support to our observations. Moreover, reciprocal BLAST from the transcriptome hits always returned a bilaterian *(L. anatina* or *S. kowalevskii)* best hit, not fungal, protist or other non-bilaterian GH13_1 sequence.

The new data unveils an evolutionary story more complicated than previously supposed. There are two alternative explanations. The first explanation is that several HGTs occurred from non-bilaterian to both hemichordate and Lophotrochozoa ancestors. The second explanation is that the ancestral GH13_1 gene was not lost in all bilaterian lineages, but remained (given the current data) in hemichordates, Brachiozoa, Bryozoa, and in scattered lineages across Mollusca and Annelida.

The hypothesis of HGT requires several such events between metazoans. It implies that HGTs obviously happened after the split of the two main branches of bilaterians, protostomes and deuterostomes, otherwise the transferred copies should have been lost in most phyla, like in the alternative hypothesis. More precisely, in the case of Lophotrochozoa, this would have occurred before the diversification of this clade and after its divergence from the Platyzoa, some 700 mya (Kumar *et al.* 2017); in the case of hemichordates, after diverging from their common ancestor with the echinoderms, and before the divergence between *S. kowalevskii* and *Ptychodera flava,* i.e. between 657 and ca. 435 mya (Kumar *et al.* 2017). Therefore, we may infer *at least* two HGTs, each early in the evolution of the phyla, with a number of subsequent losses in Lophotrochozoa (Fig. 3A). The donor species, given the sequence clustering in the trees, could be related to cnidarians. However, we have underlined that the intron-exon structures of the bilaterian sequences were most similar to the one of the sponge, and that the cnidarian GH13_1 amylases had very different structures. This may be possible if the donors were related to cnidarians, perhaps an extinct phylum or an ancestor of extant Cnidaria, but had conserved the ancestral structures exemplified by the sponge and the placozoan. Indeed, if the structure shared by the sponge, the placozoan and the bilaterians reflects the ancestral state, cnidarians must have undergone a drastic rearrangement of the intron-exon structure of this gene. This would be in line with the long internal branch leading to this clade in the trees (Fig. 1), which suggests accelerated evolution.

**Figure 3:**
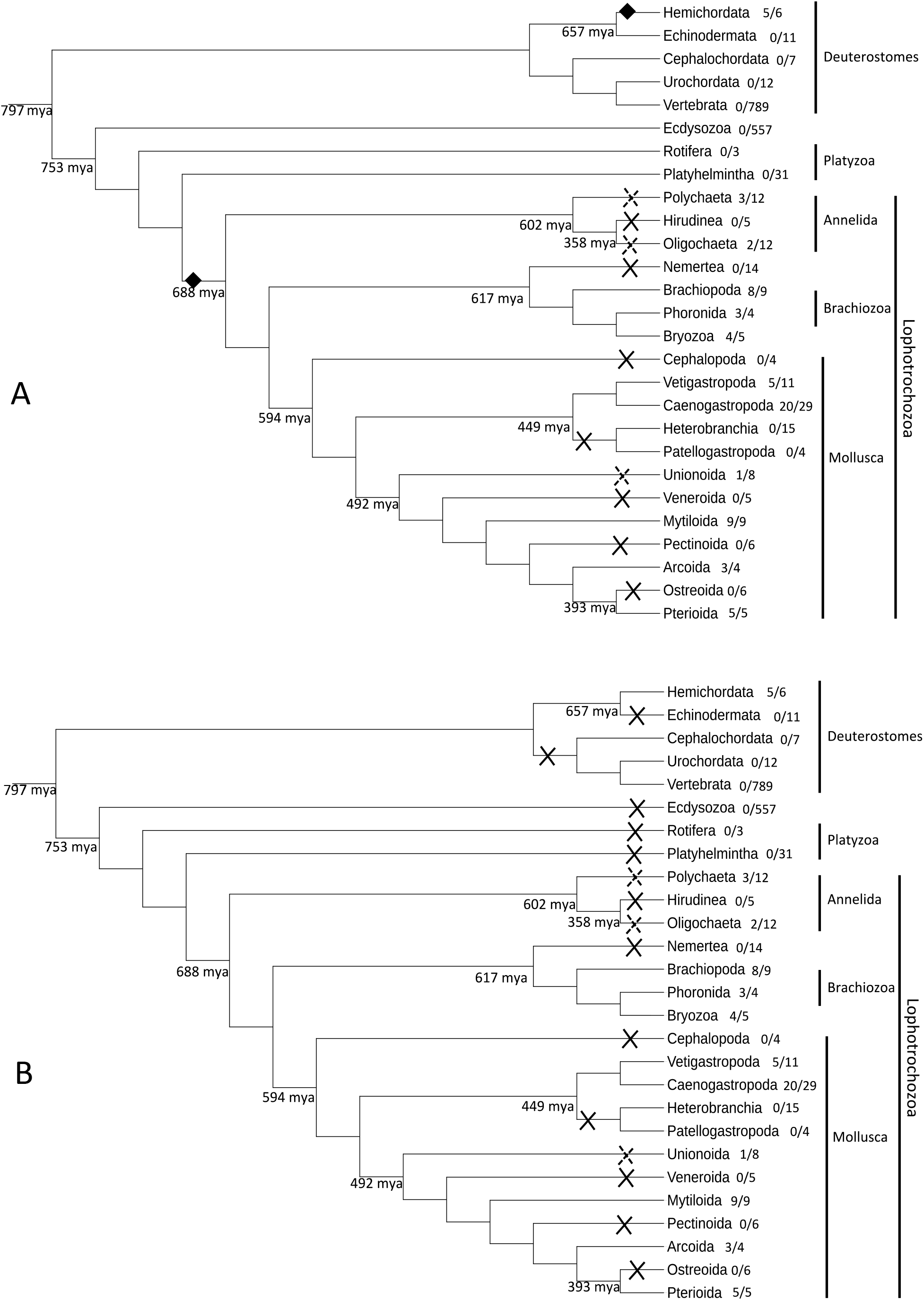
Two scenarii of HGT/gene losses of the GH13_1 genes. HGT or gene loss events were plotted on one of the proposed phylogenies of Bilateria, adapted from Plazzi *et al.* (2011); Kocot (2015); Kocot *et al.* (2017); Luo *et al.* (2015); Luo *et al.* (2018); Uribe *et al.* (2016). Fractions after the lineage names are the number of species showing GH13_1 sequences over the total number of species investigated. A: HGT hypothesis. Black diamonds represent the HGT events, crosses indicate subsequent GH13_1 loss events. B: Gene loss hypothesis. Crosses indicate GH13_1 loss events. Dashed crosses indicate lineages for which only a fraction of the available reliable genome or transcriptome data were found to contain a GH13_1 sequence. Divergence times are from Kumar *et al.* (2017).

The alternative hypothesis of massive GH13_1 gene loss in most phyla except the ones where we found such sequences seems no more parsimonious. It requires many losses, the number of which depends on the phylogeny used. For instance, considering the phylogeny shown in Fig. 3B, regarding deuterostomes, one loss occurred in echinoderms and another one in chordates. In protostomes, one GH13_1 loss in ecdysozoans, and independent losses in Platyzoa and in several lophotrochozoan lineages would be required to produce the observed pattern.

However, although not parsimonious in terms of number of events, we would rather favor the gene loss hypothesis, because this is a common phenomenon, especially given how ubiquitous co-option is (Flores and Livingstone 2017; Hejnol and Martindale 2008). In this respect, the GH13_15/24 gene that was acquired from a bacterium is a type of horizontal transfer akin to what Husnik and McCutcheon called a “maintenance transfer” since it allowed the original function to be maintained while the primitive GH13_1 gene became free to evolve or even to be lost (Husnik and McCutcheon 2018) (see also Da Lage *et al.* 2013). In contrast, while numerous cases of HGT from bacteria to metazoans, or from fungi to metazoans have been reported (e.g. Wybouv *et al.* 2016; Dunning Hotopp 2011, 2018; Haegeman *et al.* 2011; Crisp *et al.* 2015; Cordaux and Gilbert 2017), very few HGT events have been inferred that involve a metazoan donor and a metazoan receiver (Rödelsperger and Sommer 2011; Graham *et al.* 2012; Gasmi *et al.* 2015). Thus, our current knowledge on HGT suggests that this type of transfer might be very rare between metazoans, and that two or more such events would be quite unlikely to explain the current taxonomic distribution of metazoan GH13_1 genes. In addition, it has been shown that a seemingly patchy gene distribution suggestive of HGT may, after more comprehensive taxon sampling, turn out to be rather due to recurrent gene losses, as discussed in Husnik and McCutcheon (2018). The conservation of the intron-exon structure across phyla, probably ancestral to the metazoans, would not be surprising (Sullivan *et al.* 2006; Srivastava *et al.* 2010; Srivastava *et al.* 2008). For instance, 82% of human introns have orthologous introns in *T. adhaerens* (Srivastava *et al.* 2008).

In the present study we used the results of BLAST searches (BLASTP and TBLASTN) as raw material using the GH13_1-like alpha-amylases found in non-bilaterian animals (Da Lage *et al.* 2007) as query sequences. The stringent threshold we have set avoids retrieving irrelevant sequences belonging to other GH13 subfamilies or even other GH families. For instance, HMM search, such as in PFAM (pfam.xfam.org), shows that the domain composition of e.g. the *Lingula anatina* sequence XP_013396432 consists in an alpha-amylase domain linked to a DUFF1966 domain (DUFF1266 is also present in several fungal proteins, including obviously the GH13_1 amylase). The alpha-amylase domain is actually present in many glycosyl hydrolase families. Interestingly, the sequences found in some molluscs do not have a complete alpha-amylase domain, because they are shorter than usual (see Results). We assumed nonetheless that all the sequences we recovered belong to the GH13_1 subfamily, due to sequence similarities, as shown by the easy sequence alignment. Further, some of them have been assigned to this subfamily in the reference database CAZy.org (see Table 1). In addition, if we add sequences from the closest subfamilies, namely GH13_2 or GH13_19 (Stam *et al.* 2006) in the alignment and in the phylogenetic tree, the putative GH13_1 and the ascertained GH13_1 remain well clustered together (not shown). It is possible that modifications of a few amino acid positions could bring a change in the substrate or catalytic activity. For instance, concerning the substrate affinity, when the genome of *L. anatina* was released, the authors hypothesized a biomineralization pathway that involves acid proteins, as found in scleractinians and molluscs (Marin *et al.* 2007; Ramos-Silva *et al.* 2013). Given the calcium binding activity of alpha-amylases (Boel *et al.* 1990; Grossman and James 1993; Svensson 1994; Pujadas and Palau 2001), the presence of both GH13_1 and GH13_15/24 subfamilies in *L. anatina* opens the possibility for the neofunctionalization of one of them in the biomineralization process. In the analyses performed by those authors, no amylase was found in the shell matrix, but this does not exclude the possibility of its presence in the pathway. Moreover, the fact that in some molluscs, the sequences are incomplete compared to the brachiopod query or to the sponge and cnidarian GH13_1 amylases, and therefore probably devoid of an amylolytic function, would add credence to another function, especially considering that they are transcribed. This conjecture requires further investigation. On the other hand, the full-size GH13_1 sequences only present in a few bilaterians could have remained true alpha-amylases with the classical function, but this would make even more enigmatic why they have been conserved, either by descent or by horizontal transfer.

## Acknowledgments

We want to thank Pedro E. Vieira for introducing AD to molecular analyses. We are grateful to Didier Casane and Emmanuelle Renard for fruitful advise and discussion and two anonymous reviewers for critical reading of the manuscript. We thank Héctor Escrivà for sharing sequence data. This work was funded by the Conselho Nacional de Desenvolvimento Científico e Tecnológico (CNPq) (process no. 141565/2017-9) to AD and regular funding of the CNRS to JLDL and CG. CG was also supported by a grant from Agence Nationale de la Recherche (ANR-15-CE32-0011-01 TransVir).

